# Establishment and equilibrium levels of deleterious mutations in large populations

**DOI:** 10.1101/318428

**Authors:** Johan W. Viljoen, J. Pieter de Villiers, Augustus J. Van Zyl, Massimo Mezzavilla, Michael S. Pepper

## Abstract

Analytical and statistical stochastic approaches are used to model and predict the dispersion of mutations through a large population. These approaches are used to quantify the magnitude of a heterozygous selective advantage of a mutation in the presence of a homozygous disadvantage. Random effects such as genetic drift are accounted for, which are likely to extinguish even highly advantageous mutations while the prevalence is still low. Dunbar’s results regarding the cognitive upper limit of the number of stable social relationships that humans can maintain are used to determine a realistic community size - a reduced local subset of the total population - from which an individual can select mates. This reduction in effective population size has a dramatic effect on the probability of establishing mutations, as well as the eventual equilibrium values that are reached in the case of mutations conferring a heterozygous selective advantage, but a homozygous disadvantage, as in the case of cystic fibrosis and sickle cell disease. The magnitude of this selective advantage can then be estimated based on observed occurrence levels of a specific mutation in a population, without requiring prior information regarding its phenotypic manifestation.

**Author summary:** Deleterious mutations such as cystic fibrosis and sickle cell anemia can disperse through human populations due to the selective advantage that it bestows on heterozygous carriers, depending on environmental conditions. As its prevalence increases, the probability of generating homozygous offspring, with its concomitant selective disadvantage, also grows until an eventual equilibrium is reached between the number of carriers and wild-type individuals. In this work computer modelling is used to combine Dunbar’s anthropological observations predicting upper bounds to the number of stable human social relationships with observed prevalence levels, to estimate the absolute magnitude of the heterozygous selective advantage bestowed by such a deleterious genetic variation, without requiring knowledge regarding the specific mechanism whereby such an advantage is manifested.

## Introduction

A computational and statistical framework was created to simulate and calculate the diffusion of monogenic variations over multiple generations through a population of organisms. Although the work was originally aimed at exploring aspects of cystic fibrosis (CF) in humans, the tool is generally applicable to the modelling of the epidemiology of mutations, where there are differences in fitness and/or survival success rates between homozygous and heterozygous individuals. This is based on the observation that heterozygous carriers of some common mutations, e.g. in the cystic fibrosis transmembrane conductance regulator (CFTR) gene, seem to have a survival/fecundity advantage compared to non-carriers [1, 2], while the homozygous state results in a definite *dis*advantage. An example of a heterozygous advantage that is environmentally dependent is malaria resistance [3, 4], bringing with it the risk of sickle-cell anemia in homozygotes. Another common example of a heterozygous advantage is the MHC complex in vertebrates [5].

The implications of such a benefit (termed ‘selective advantage’) was investigated by Haldane [6], who focused on the mathematical probability of purely beneficial mutations becoming established in a population of size *N*. Haldane’s results were subsequently refined by Wright [7], Kimura [8], and others. Wright also introduced the notion of ‘effective population size’ (*N*_e_), to account for effects such as non-random mating, inbreeding and unequal sex ratios, which may influence the effectiveness of natural selection forces.

For human populations we further extend this approach by drawing on the results of Dunbar [9] and Lehmann [10] to propose a realistic range and upper limit to the size of the group from which an individual is likely to select a mate. This is analogous to the ‘breeding unit’ introduced by Wright in 1946 [11], which he termed *N*_n_, being the spatially closest individuals in a circular area with a radius of 2*σ*:

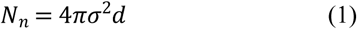

with *σ* the standard deviation of a spatially distributed 2-dimensional normal distribution around an individual and *d* the distribution density. This reflects the observation that the parents of an individual organism are more likely to be proximate than remote, and, as Nunney suggests [12], that *N*_n_ will be relatively constant, subject to the assumptions that *σ* is a property of the species and that there is a negative correlation between dispersal and density, i.e. that one can normalize for *d*. This leads to a parental probability distribution solely dependent on distance *r*:

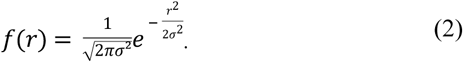

In the case of human populations, we propose that the concept of distance (*r*) should be reinterpreted as social proximity, rather than necessarily physical proximity, due to the global mobility (i.e. potentially high dispersal) that has been attained by humans in recent times, and which does not necessarily translate into an increase in *N*_n_ (which is also the pool from which an individual would usually select a mate). Based on primate studies and the size of the human neocortex, Dunbar posits a nominal maximum group size of 148 for humans (usually rounded up to 150), with a 95% confidence interval between 100 and 230 [9], but also stresses that this is an upper value, which is only approached under extreme environmental stress, where the significant time and energy investments of maintaining close social bonds are repaid by the survival benefit realized by being part of a larger group. In more prosperous times, group sizes tend to reduce.

This aspect is one of the major novelties of our framework, that allows us to simulate large populations while modifying the community size (*N*_n_) in order to create scenarios to study how local selection affects the pattern of deleterious variants.

We then explore the effect of this community size, in conjunction with the selective advantage of a given mutation, on the probability that such a mutation will become established in the population, and on the eventual equilibrium levels that are reached, especially in the case of mutations that are beneficial when heterozygous, but pathogenic (or less beneficial) when homozygous. Such mutations do not necessarily become ubiquitous once established. Conversely, if we know the occurrence levels of such a mutation in a population, we can use our model to estimate the selective advantage that it confers on heterozygous carriers, without requiring any knowledge of the specific manifestation and mechanism of such a selective advantage.

## Results

### Establishment of Mutations

For haploids, the selective advantage *s* conferred by a specific mutation is defined as an additive term, such that carriers would, on average, have (1+*s*) times as many offspring as wild-type (i.e. non-carrier) individuals [13]. Note that *s*, normally termed a ‘selective advantage’, can also be a negative number, implying a disadvantage by resulting in fewer offspring of mutant carriers, whether through lowered fertility, or decreased survival to procreative age (which in some sense is the same thing - irrespective of the mechanism, the result being a reduction in the number of offspring compared to wild-type individuals). Although the case of *s <* 0 has the physical interpretation of a selective *disadvantage* (purifying selection), the case of *s* < −1 is meaningless, and as such the parameter *s* should be constrained on the interval [-1,∞).

Using a deterministic model, any beneficial mutation (that is, with *s* > 0) will inevitably grow in prevalence, guaranteeing eventual fixation in the population. However, in reality genetic drift causes random fluctuations in the frequency of lineages, which can easily extinguish even highly beneficial mutations when its prevalence is low, as will be shown. A stochastic treatment is used to analyze such situations, which especially apply whenever a new mutation appears *de novo* in a single individual. The mutation will only be established in the population (and only then a deterministic model may be applicable) if this mutation survives genetic drift. During admixture between different populations, ‘new’ alleles are introduced at significant levels into both groups. Under such conditions (relatively large populations and high prevalence) the alleles could be considered to already be established and therefore to be less subject to the vagaries of genetic drift, but rather with their eventual fate dominated by the relative selection coefficients that the alleles confer (i.e. closer to the deterministic case).

Addressing the fixation probability *π* of a single copy of an advantageous allele in a large population, Haldane [6] found that

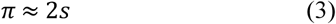

if *s* is small. Negative or zero values of *s* always lead to eventual extinction.

### Stochastic Determination of Fixation Probability

To investigate the statistical fate of a dominant mutation with selective advantage *s* (for both heterozygous and homozygous cases) in a large population, our simulation tool was configured to introduce a single mutation in a virtual population (population size *N* = 2.5×10^5^), and then to cycle through the generations until the mutation either becomes ubiquitous or extinct. This was repeated 200000 times for each point in the graph shown in **Error! Reference source not found.**, for a total of ~112 million generations. The results confirm Haldane’s prediction, including his *caveat* that it is only valid for small values of *s*. The stochastic simulation initially closely follows the *π* = 2*s* line, but then gradually deviates from it as *s* increases. This is expected: the fixation probability cannot exceed 1; it can only approach unity asymptotically as *s* grows.

### The Price of Success

#### Homozygosity

A mutation that is purely beneficial will completely displace the wild-type allele only if it successfully runs the gauntlet of genetic drift while still rare. In a diploid population, however, heterozygous carriers may derive a selective advantage from a given mutation, while homozygosity results in a reduced selective advantage (or even disadvantage), such as in the case of CF, sickle-cell disease and others. As the prevalence of such a mutation in a population increases, the probability of producing homozygous offspring also rises, to the point where the relative disadvantage of homozygosity exactly balances the heterozygous advantage. An equilibrium is reached, depending on the relative magnitudes of the effects, as well as the population parameters, especially the effective population size *N*_e_ and the neighborhood size *N*_n_.

#### Environmental Factors

The striking geographic correlation between the distribution of the sickle-cell allele and the prevalence of malaria [3] demonstrates this effect. As long as the local population is exposed to malaria, the mutation (if present) confers an advantage (*s* > 0) to heterozygous carriers, and rises to prevalence levels limited by the negative effects caused by the associated increase in homozygous individuals. Where malaria is absent, there is no selective advantage (*s* may even be slightly negative), and the allele becomes extinct. Environmental variables definitely matter.

#### Migration, Selection and Inbreeding

A shortcoming of Haldane’s approach is that the population *N* is assumed to be large, constant and with equal sex ratios and random mating. This is not normally the case. Many subsequent researchers have addressed this [7, 13, 14] and have introduced the concept of an *effective population size N*_e_ which would result in the same variance as the actual population in question. Usually the effective population size is smaller than the census size (*N*_e_ < *N*), and there is the even smaller community size (or neighborhood number) *N*_n_ << *N*_e_ which affects genetic differentiation between subpopulations: the smaller *N*_n_ the larger the differentiation between them, due to the decreased dispersal distance and increased genetic drift (considering limited or no gene flow) [12].

### Probabilistic Analysis

This section presents a heuristic deduction of a simple relationship between the long-term prevalence of the mutation and the heterozygous advantage proposed by the model in this paper, assuming a large two-dimensional neighborhood size. The (random) state of the individual at grid point (i,j) in the population array can be described as a stochastic process 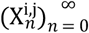, where *n* denotes the number of generations from the beginning. We use *w* for the wild-type, *e* for the heterozygous and *m* for the homozygous states respectively.

The states of different individuals are distributed identically, in other words they are statistically indistinguishable, because the development of a specific individual depends only on its neighbors and not the absolute position (*i*, *j*). This is due to the fact that the population is closed upon itself, so that there are no edges or other position-specific effects. Hence we can drop the superscripts *i* and *j* and denote the state of a generic individual at the *n^th^* generation by *X_n_*.

Our concern is with the temporal evolution of the quantity *p_n_*: = *P*(*X_n_* = *e*); that is, the probability that a generic individual is a heterozygous carrier of the mutation. Additionally, we would like to derive the existence and value of the equilibrium level *p* so that *p_n_ = p* implies *p_n_*_+1_ = *p*. For large populations, the value of *p* indicates the prevalence of the mutation, that is, the percentage level at which the heterozygous advantage balances out the homozygous disadvantage, on average.

We assume, for ease of display, that the homozygous *dis*advantage is 100% (i.e. *s_hom_* = −1) so that homozygous individuals produce no offspring. We also assume that mutations only occur once, at generation *n*=0.

We use the following notation:

*F_k_*: The event that *k* of the *N* neighbors are in state *e*.

*E*_10_: The event that the first parent of the individual is in state *e*, but not the second. (This event depends on the generation *n* but for readability we omit *n* from the notation.)

*E*_01_: The event that the second parent of the individual is in state *e*, but not the first.

*E*_11_: The event that both parents of the individual are in state *e.*

*E*_00_: The event that neither of the parents of the individual are in state *e*.

*s*: The heterozygous selective advantage (*s_het_*).

Before we deduce the difference equation for *(p_n_)*, we first compute, for each specific value of *k*, the probability that an individual is heterozygous in the next generation given *k* currently heterozygous neighbors. Since for each individual exactly one of *E*_00_, *E*_01_, *E*_10_ or *E*_11_ must happen, and expanding the conjunction of events into conditional probabilities, we get

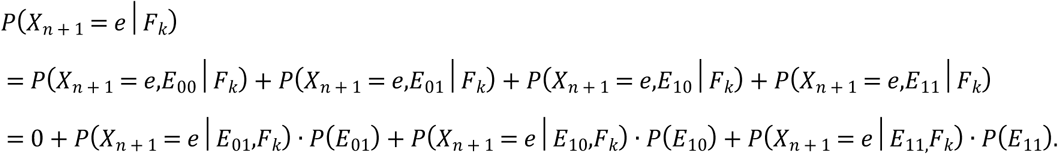

Using Mendelian inheritance probabilities we have 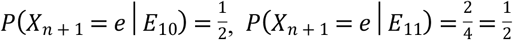 and so on. Thus

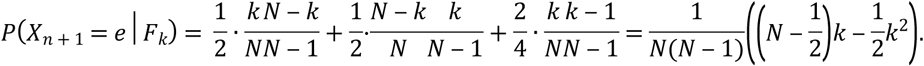

So for each value of *k* we have the probability that *X_n_*_+1_ is heterozygous given that *k* of the parents in its community are heterozygous. We can combine these conditional probabilities by conditioning on the number of neighboring parents that are heterozygous:

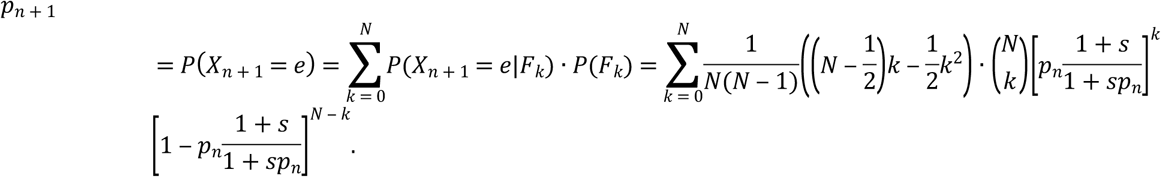

From the first and second moments of the binomial distribution we know that for any *q*, 0 ≤ *q* ≤ 1, we have

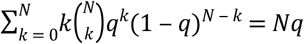
and

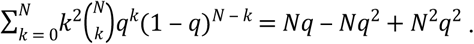

Applying these identities with 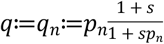, our equation simplifies to

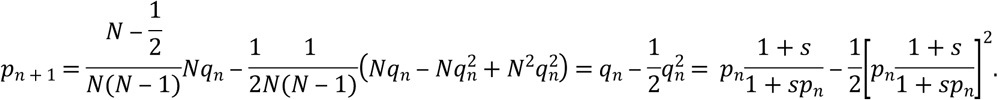

Equilibrium will be achieved where *p* = *p_n_*_+1_ = *p_n_*. This leads to a quadratic equation which can be solved easily, although the resulting expression is somewhat cumbersome.

In the case where both *s* and *p_n_* are small, the product *sp_n_* is a second-order term, so a first-order approximation to the equilibrium value is given by

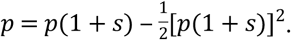

Elementary algebraic operations give

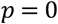

or 

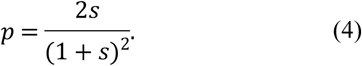

As before we now compare our analytic equilibrium prediction in Equation (4) with our stochastic simulation (setting *N*_n_ to a large value, *s_hom_* = −1, and varying the heterozygous selection advantage *s_het_*), achieving satisfactory correspondence as shown in Fig 2, where for small *s* (and resultant small *p*) the results are essentially identical, as expected.

**Fig 1.**
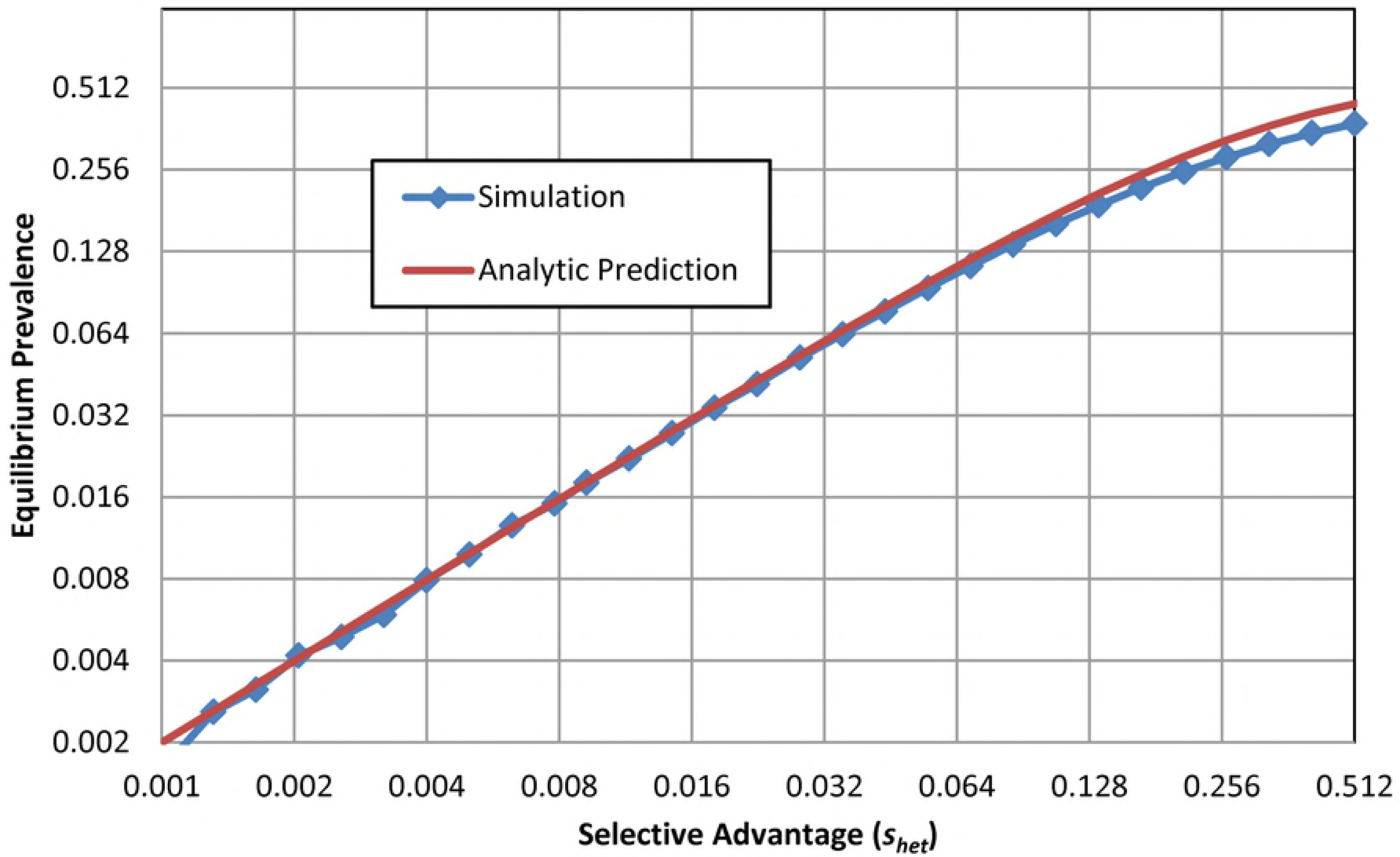
Establishment probability of a beneficial mutation. Fixation rate for a single mutation as a function of (positive) selective advantage (*N*=*N*_n_= *N*=2.5×10^5^).

**Fig 2.**
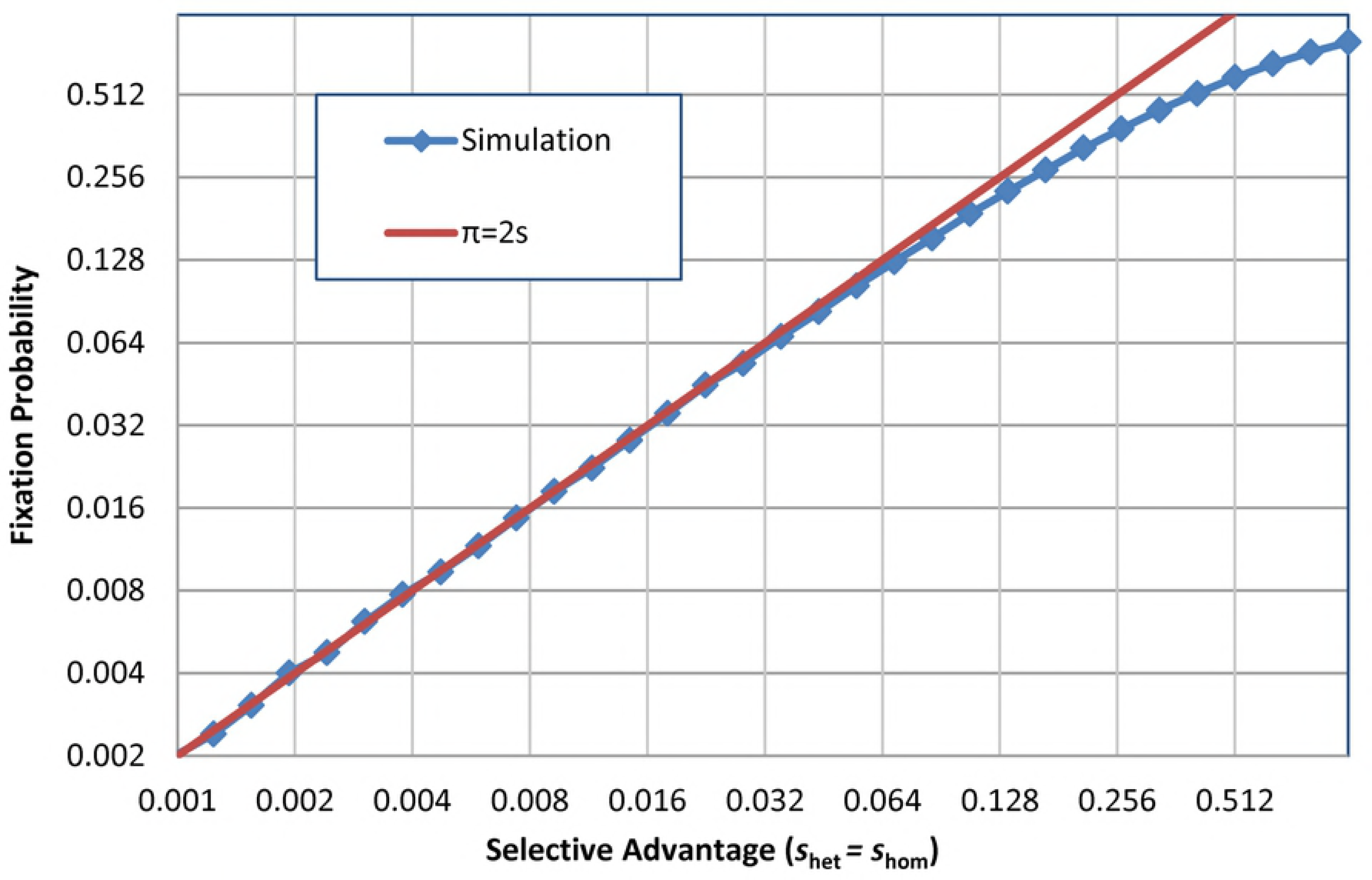
Equilibrium levels of a deleterious mutation. Equilibrium level as a function of (positive) heterozygous selective advantage, with homozygous selective advantage = −1. (*N*=*N*_n_=9×10^6^, *s_hom_*=-1, generations=5000.)

## Numeric Simulation Results

The computational tool described above was used to explore the behavior of different types of monogenic variation under various conditions.

### Single Runs

#### Homozygous deleterious

**Error! Reference source not found.** was generated to test the numerical simulation, using two different selective advantage values *s* for heterozygous carriers, with homozygous carriers having a 100% disadvantage (i.e. none survive to procreate, which approximates reality for CF). To stabilize at a prevalence of 4% (reflecting the approximate CF carrier frequency in the European population) requires either a large community size *N*_n_ and *s* = 2.132%, or, more realistically, a community size of 150 (approximately Dunbar’s number for humans) and *s* = 2.95%. Both of these scenarios eventually reach equilibrium at 4%, after hundreds of generations.

The implication is that the actual advantage of being a carrier is indeed higher than the result that would be obtained without taking into account the limited community size, where consanguinity augments the probability of producing homozygous offspring.

#### Positive selection

Lactase persistence is associated with the LCT gene (MIM 603202) and is especially prevalent in European populations, with evidence of strong recent selection during the last 5000-1000 years [15], coinciding with the domestication of cattle and a rise in dairy farming. In such a setting, the ability of adults to derive nutrition from dairy confers an obvious advantage.

Bersaglieri *et al* [15] estimate the selection coefficient for lactase persistence to be between 0.09 and 0.19 for the Scandinavian population (where the prevalence of the −13910T allele currently exceeds 80%). In Fig 4 a conservative value of 0.1 (10%) was assumed as the selective advantage for both heterozygotes and homozygotes. Starting from an initial low base, the prevalence of heterozygotes grows to a maximum value of just over 30%, by which time it has already been surpassed by homozygotes, which asymptotically approach 100%, given enough time (generations). After just more than 200 generations, 80% of the population carries the mutation – at a nominal 29 years per generation [16] this corresponds to 5800 years, which is near the lower end of the antiquity estimate for dairy farming. However, it seems reasonable to assume that dairy farming also took a long time to become common, which implies that the availability of milk, and hence the effective selective advantage of LCT, gradually increased to its current value.

**Fig 3.**
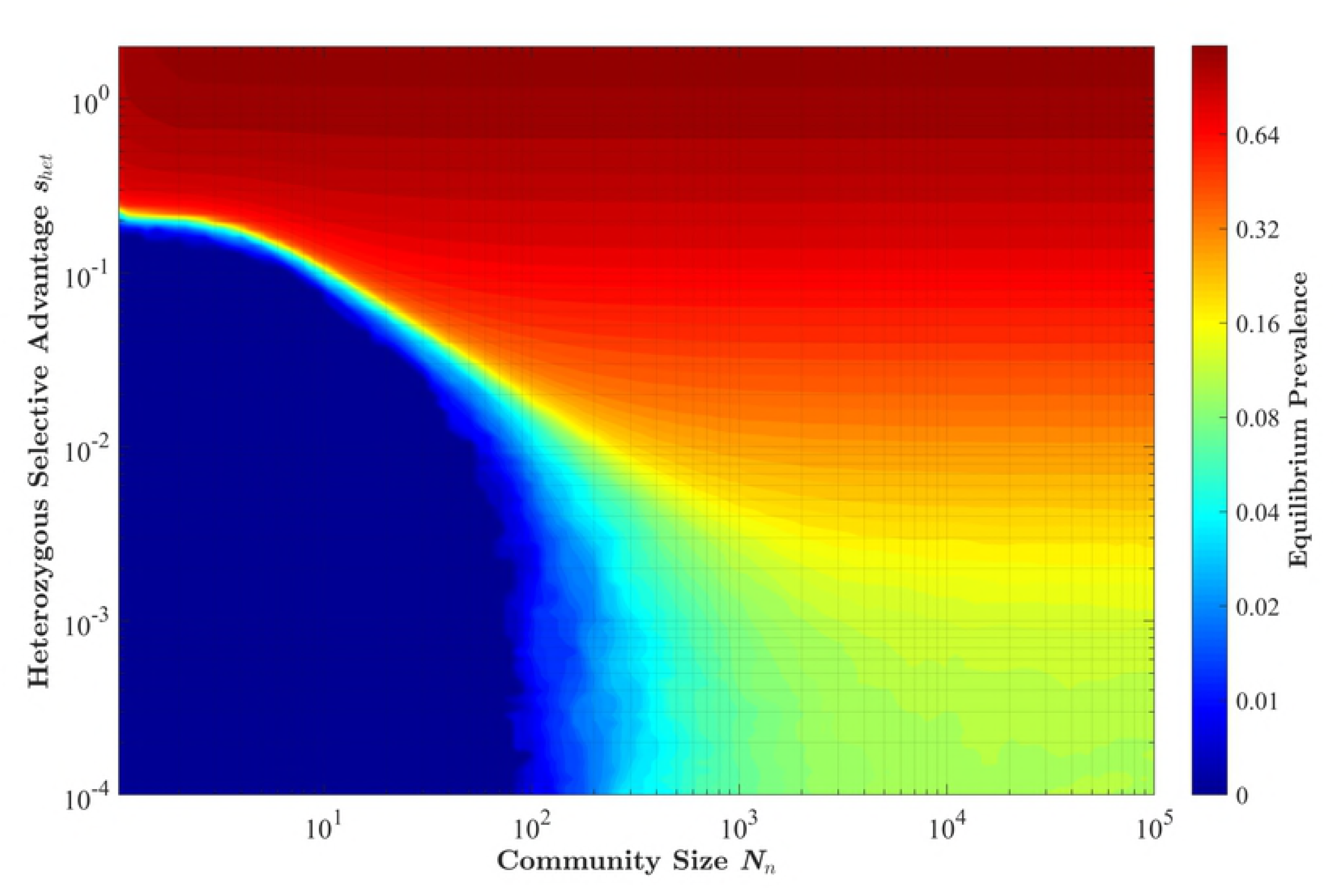
CFTR Carrier prevalence. Results for numerical simulations of mutation dissemination in a test case to match the ~4% CFTR mutation carrier frequency in the European population. *N*=10^8^, with each curve the average of four independent simulation runs to 3000 generations.

**Fig 4.**
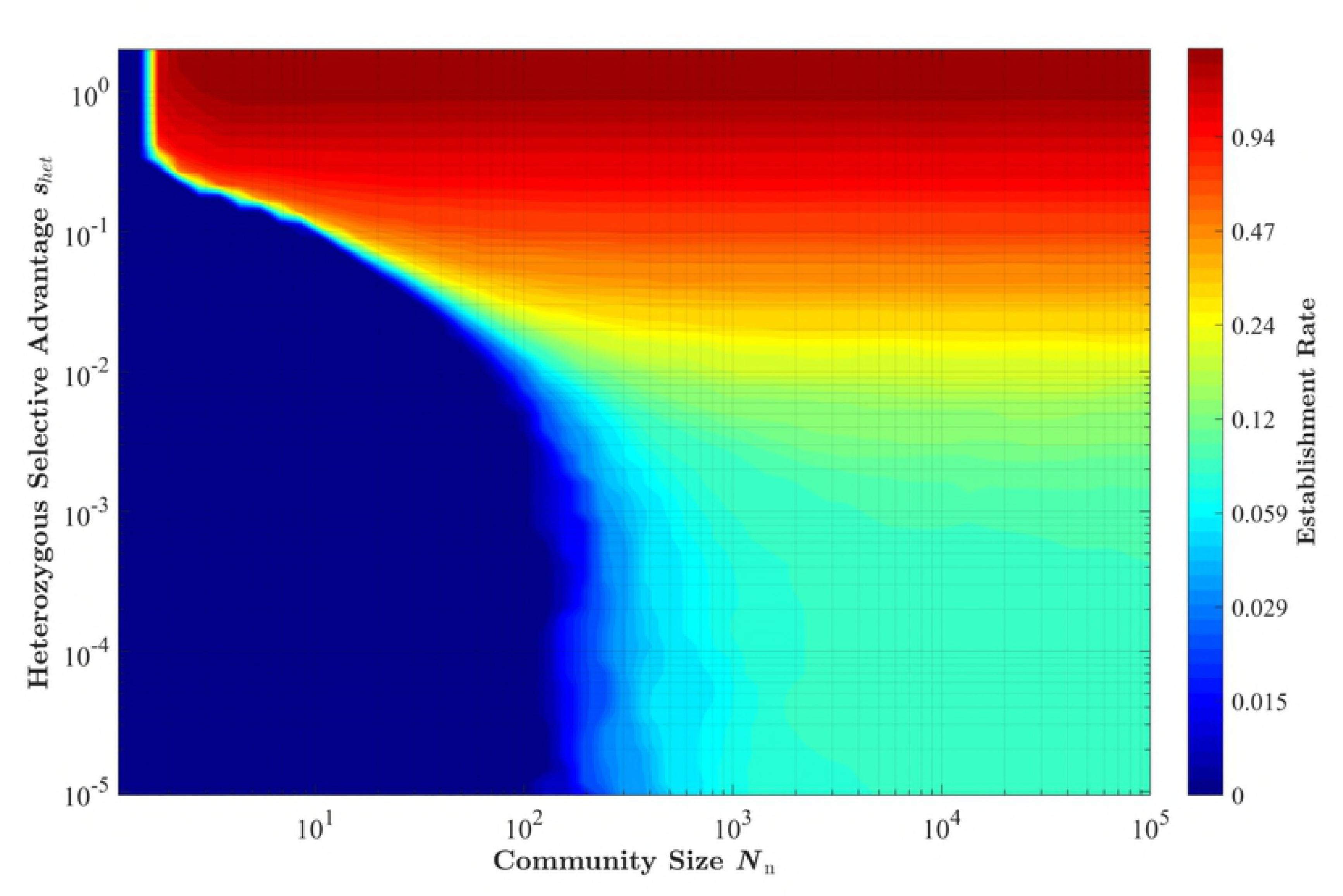
Lactase persistence. The spread of a purely beneficial mutation (*s_het_*=*s_hom_*=0.1, *N*=5×10^6^, *N*_n_=150).

#### Monte Carlo Analyses

A Monte Carlo analysis was performed, whereby multiple runs were executed automatically, while varying the starting conditions and input parameters. This enabled the compilation of statistical results over millions of trials.

Fig 5 illustrates how the eventual equilibrium prevalence of a mutation (with homozygous selection coefficient *s_hom_* = −1) depends on both the community size and the heterozygous advantage that it confers on a carrier.

**Fig 5.**
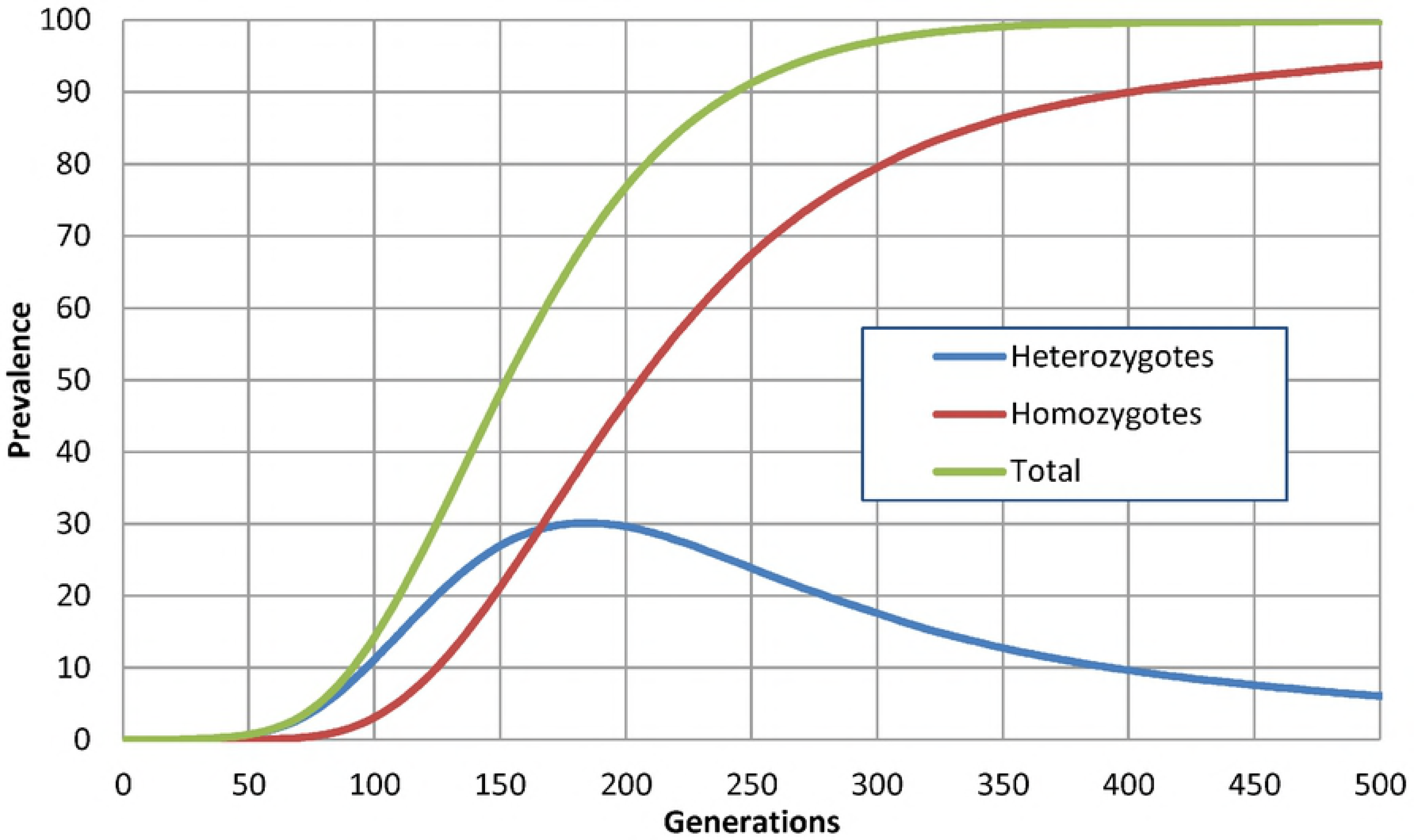
Equilibrium prevalence percentage as a function of Community Size and Heterozygous Selective Advantage. Monte Carlo run results (*N*=10^6^, *s_hom_* = −1, max 10000 generations per point).

Fig 6 shows the probability that a single mutation will indeed become established, once again as a function of community size and the heterozygous selective advantage that it confers, while keeping the homozygous selection coefficient equal to −1, which approximates the case found in diseases such as CF. When the community size is large, the behavior approaches that seen in **Error! Reference source not found.**. From the data generated by the simulations underlying this plot one can also extract statistics regarding the average survival (in generations until extinction, if this happened) and maximum prevalence that a given mutation attained.

**Fig 6.**
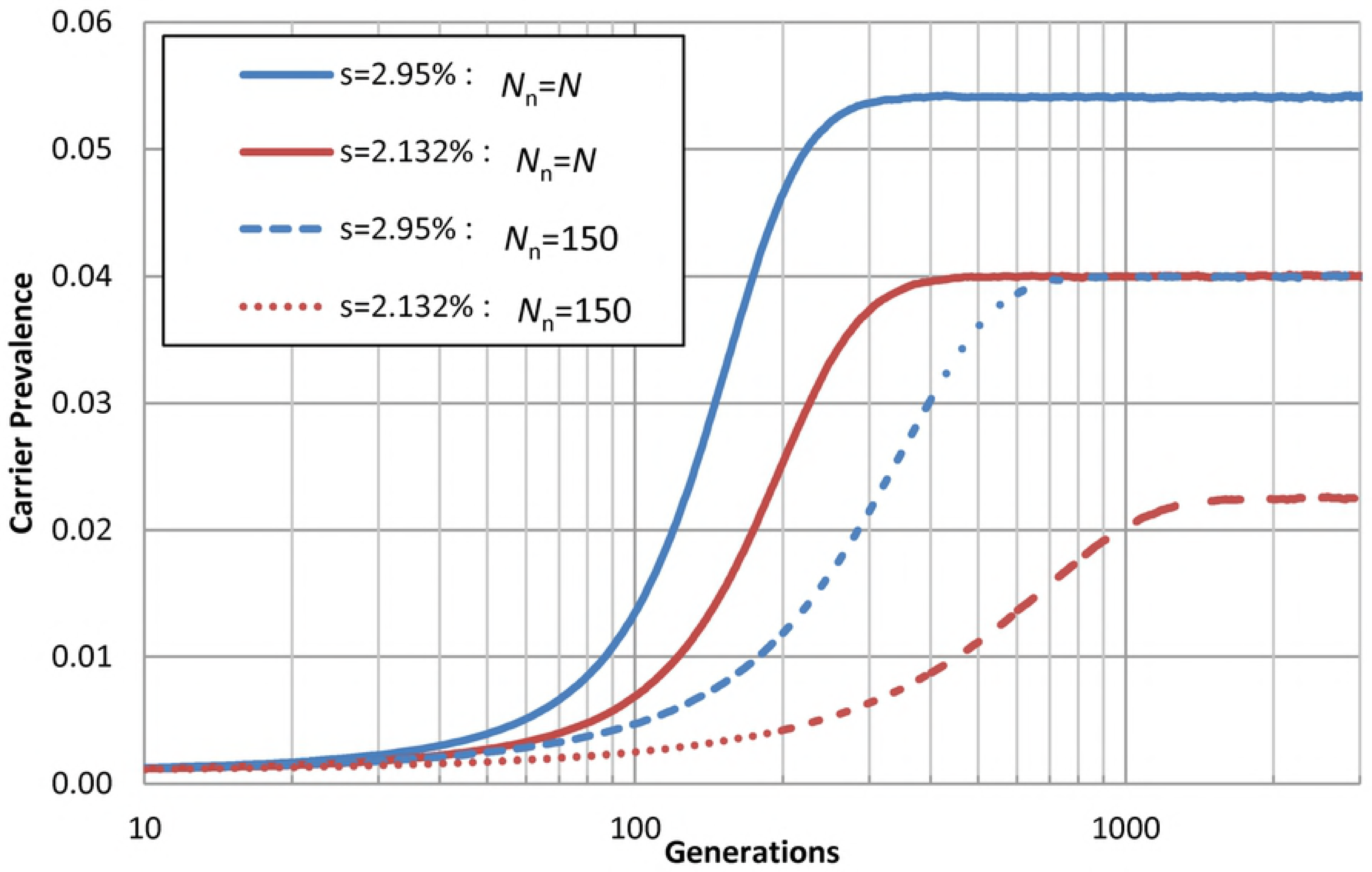
Rate of establishment as a function of community size and heterozygous selective advantage. Monte Carlo run results (*N*=10^6^, *s_hom_* = −1).

### Discussion

1. The close correlation of the simulation results with our own theoretical derivations, as well as with theoretical predictions in the literature, inspires confidence in the model that we have created.
2. By using actual prevalence values for a given monogenic variation (such as for CF), and assuming that equilibrium has been reached, one can estimate the selection coefficient, without requiring any knowledge of its specific manifestation and mechanism.
3. Assuming equilibrium, we estimate that being a heterozygous CFTR mutation carrier confers a selective advantage of at least 3% (depending on our assumption regarding *N*_n_).
4. Deleterious recessive mutations as considered here *must* confer some survival/fecundity advantage, otherwise they never reach noticeable prevalence levels.
5. Most mutations, even significantly advantageous ones, do not become prevalent.
6. The probability of a mutation becoming prevalent, as well as the equilibrium level it eventually attains, is strongly linked to the combination of selection coefficient (which includes environment) and the effective community size.
7. The low probability of even an advantageous mutation becoming prevalent, the number of generations it takes to spread through a large population, and the large number (2000+) of known CFTR mutations in humans seem to indicate that the *de novo* mutation rate may be surprisingly high. In addition, from the ExAC database [17] CFTR genes seem to harbor a significantly greater number of missense variations with respect to the expected (expected n. missense=418.8, observed n. missense=671, missense z-score –6.03).
8. The simulation framework allows us to create scenarios to study selection and in particular balancing selection, introducing the concept of community size, which could be useful to simulate a realistic scenario of population structure and isolation inside larger populations.

## Materials and Methods

A numeric simulation tool was created to facilitate the stochastic exploration of the fate of monogenic variations.

### Assumptions

1. Only a single gene (with a mutation/variation that changes the procreation probability of carriers) is considered.
2. Compared to the general (non-carrier) population, heterozygous and homozygous carriers of mutated genes can have different survival/fecundity rates.
3. Individuals select mates for procreation purposes from a limited community *N*_n_, which in general is much smaller than the size of the entire population. For a population of humans, Dunbar and Lehman motivate an upper cognitive limit on the number of people with whom an individual can maintain stable social relationships [9, 10]. This is used to inform the realistic community size from which an individual can select a mate, in this model. Dunbar’s number for humans is estimated to lie in the range of 100 to 230, with 148 being the nominal value. Values for *N*_n_ of this size and smaller are considered to be reasonable for human populations.
4. Constant environmental conditions are assumed across the entire population – although provision is being made for different geographical conditions to reflect situations such as sickle cell disease and malaria as discussed above - and all simulations were conducted with this assumption of constant conditions in mind.
5. At present the effective population number *N*_e_ is assumed to be large (comparable to the total population size *N)*, and specifically much larger than *N*_n_, i.e. *N*_e_ >> *N*_n_. This is adjustable, however, to allow for exploration of effects in smaller populations and genetic isolates.

### Data Structure

1. A two-dimensional array is created, with every element representing an individual in the population. Computational resources now make it feasible to create and process simulated populations consisting of millions, and even billions, of individuals.
2. Each individual has one of four possible states: dead, no mutation, heterozygous, homozygous.
3. The population array is closed upon itself, with edges wrapping around. This eliminates any edge effects that discontinuities may otherwise introduce.
4. Physical proximity in the elements of the population array is used as a proxy for social closeness – i.e. an individual is more likely to breed with another nearby individual than with a remote one, according to a two-dimensional normal (Gaussian) probability distribution.
5. The effective size of the community around each individual is changed by varying the standard deviation of the normal distribution, with *N*_n_ as in Equation (1).

### Simulation Procedure

For a given set of parameters, the following steps are executed:

1. Set population size, community size *N*_n_, initial carrier prevalence, advantage/disadvantage for carriers, and *de novo* mutation probability.
2. Initialize the population randomly with desired initial fraction of carriers.
3. For each individual in the population, change its status to dead with a probability dependent on its current status, to statistically reflect the advantage/disadvantage associated with its status.
4. For each element of the population array:

a. Randomly select two distinct non-dead parents from its community *N*_n_, according to the proximity probability distribution as in Equation (2).
b. Generate a status according to Mendelian inheritance probabilities from the two parents.
c. Randomly introduce a *de novo* mutation with a specified (typically low and possibly even zero) probability.
5. Update population statistics and display.
6. Return to step 3.

### Code Availability

The simulation tool was developed in Borland Delphi. Source code and the binary executable is available at https://github.com/JohanViljoen/PopSim.

## Supporting information Legends

**S1 File. Raw data for Figs 1 - 4.** *Tab 1*: An Excel spreadsheet containing the numeric simulation results compared with Haldane’s prediction for the fixation probability at varying selective advantage. *Tab 2:* Numeric simulation results compared with our analytically derived prediction for the equilibrium prevalence of a mutation that is homozygous deleterious (*s* = −1) but with *s* > 0 (i.e. positive selective advantage) when heterozygous. *Tab 3:* The numeric simulation results four different combinations of community size *N*_n_ and *s*, showing the different establishment rates and eventual equilibrium levels. *Tab 4:* The numeric simulation results demonstrating the spread of lactose persistence at a selective advantage of 0.1 (10%) for both heterozygous and homozygous individuals, using a community size *N*_n_ = 150.

**S2 File. Raw data for Fig 5.** A text file containing the summarized results of 43 Monte Carlo runs comprising 305554892 generations (population size 10^6^) which took a total of 256 days, 17 hours and 53 minutes of processor core time to complete, spread between 6 PCs with 2-6 cores each.

**S3 File. Raw data for Fig 6.** A text file containing the summarized result of 172 Monte Carlo runs comprising 600948539 generations (population size 10^6^) which took a total of 600 days, 11 hours and 39 minutes of processor core time to complete, spread between 6 PCs with 2-6 cores each.

